# *Cyclocarya paliurus* polysaccharides improve high fat diet-fed mice lipid dysmetabolism through gut microbiota

**DOI:** 10.1101/2024.03.30.587420

**Authors:** Chengyan Huang, Yanmei Jia, Ping Chai, Huizhen He, Wei Liu, Lirong Cheng

## Abstract

Obesity is associated with low-grade chronic inflammation and intestinal dysbiosis. *Cyclocarya paliurus* (C. *paliurus*) has been shown to improve lipid metabolism and gut microbiota communities in animals and humans; however, it remains to know whether *Cyclocarya paliurus* polysaccharides (CCPP) prevents obesity through gut microbiota. Here, we found that high fat diet promoted the lipid accumulation and intestinal microbiota dysbiosis in mice, while drink *Cyclocarya paliurus* polysaccharides reduces body weight, inflammation, alleviated the lipid accumulation, and reversed gut microbiota dysbiosis, including the diversity of intestinal microbiota, relative abundances of *Lachnospiraceae*. By transplanting faeces from CP-treated mice to HFD-fed mice, the effects of anti-obesity and microbiome regulation can be transferred through gut microbiota. Our results indicate that *Cyclocarya paliurus* polysaccharides can prevent gut dysbiosis and obesity-related metabolic disorders in obese individuals, the mechanism is related to the remodeling of intestinal flora.

## 1. Introduction

Obesity has become one of the public health problems in the world. According to the World Health Organization (WHO), in 2016, more than 1.9 billion adults aged 18 and over were overweight, of whom more than 650 million were obese. Obesity rates around the world have almost tripled since 1975(1). Obesity is an important factor in inducing a variety of diseases, including hypercholesterolemia, hypertriglyceridemia, type 2 diabetes, cardiovascular disease and multiple cancers(2–4). The prevention of obesity has become among the most challenging concerns for modern society. There is growing evidence that the gut microbiota may serve as an important modulator of obesity by affecting the absorption of nutrients in the intestine(5, 6).

Gut microbiota is a large and diverse group of microorganisms living in the human gastrointestinal tract. The gut microbes themselves and their metabolites can regulate health and play an important role as a bridge between diet and host(7). The change of the gut microbiota associated with the occurrence of some diseases, such as obesity(8), gastrointestinal diseases(9), type 2 diabetes(10), and non-alcoholic fatty liver disease(11). A study indicates that changes in the composition of the gut microbiota are associated with the development of obesity and its associated metabolic disorders(12). An increased ratio of the major phyla *Firmicutes*/ *Bacteroidetes* and changes in several bacterial species can promote the development of obesity in both dietary and genetic models of obesity in mice(13, 14). Some plant extracts have been found to reshape intestinal flora and improve obesity induced by a high-fat diet(6, 15, 16), providing a new way to prevent obesity.

LPS is a powerful pro-inflammatory molecule from the cell wall of Gram-negative bacteria. It promotes the synthesis and release of inflammatory factors tumor necrosis factor alpha (TNF-α), interleukin-1β (IL-1β), and interleukin-6 (IL-6) through activating TLR-4 and downstream NF-kB and JNK inflammatory pathways(17).

Chronic low-grade inflammation (CLGI) is central to the pathogenesis of obesity(18, 19). It damages pancreatic beta cells, disrupts insulin action, and mediates glucose intolerance in obesity(20). Systemic CLGI is identified by elevated circulating levels of inflammatory cytokines, such as TNF-α, IL-1β, and IL-6, which act as molecular mediators and are responsible for the progression of the response to a systemic level encompassing multiple organs(19).

C. *paliurus*, belonging to the genus *Cyclocarya* (Juglandaceae) and commonly called the “sweet tea tree”, is only grown in southern China(21). The leaves of C. *paliurus* have been widely used in traditional Chinese medicine for a long time to treat T2DM(22, 23). Recently, many experimental studies have revealed that *C. paliurus* and the extractions of C. *paliurus* have the abilities to reduce blood glucose and lipid(23–25). *Cyclocarya paliurus* polysaccharide (CCPP), a primary active component in the leaves of *C. paliurus*, has found to possess many bioactivities(26), such as hypolipemic(27),anti-inflammatory(28),immunomodulatory and anti-cancer(29), as well as its ability to treat hyperglycemia(21, 23).

Recent studies have found that the hypoglycemic action mechanism of CCPP is also closely related to its ability to regulate host nutrition and energy metabolism by regulating intestinal microbiota(30). However, the relationship between the hypoglycemic effect of CCPP and intestinal flora still needs to be explored. Here, we studied whether the gut microbiota improved by C. *paliurus* could play a hypoglycemic role, laying a further foundation for the improvement of the relationship between energy metabolism and intestinal flora by C. *paliurus*.

## 2. Materials and methods

### 2.1 Animal and diet

All animal experiments reported in this manuscript were approved by the Animal Care and Use Committee of Institute of Fenyang Collage of Shanxi Medical University (approval no.2023022). C57BL/J male mice were purchased from Shanxi Medical University Laboratory Animal Central (Taiyuan, China), permission code SCXK2019-0004. All animals had free access to food and drinking water and housed in an independently ventilated cages during the experimentation. After experiment, all animals were euthanized. The formula and dosage of normal food and high-fat diet food are shown in Table 1.

**Table1:**
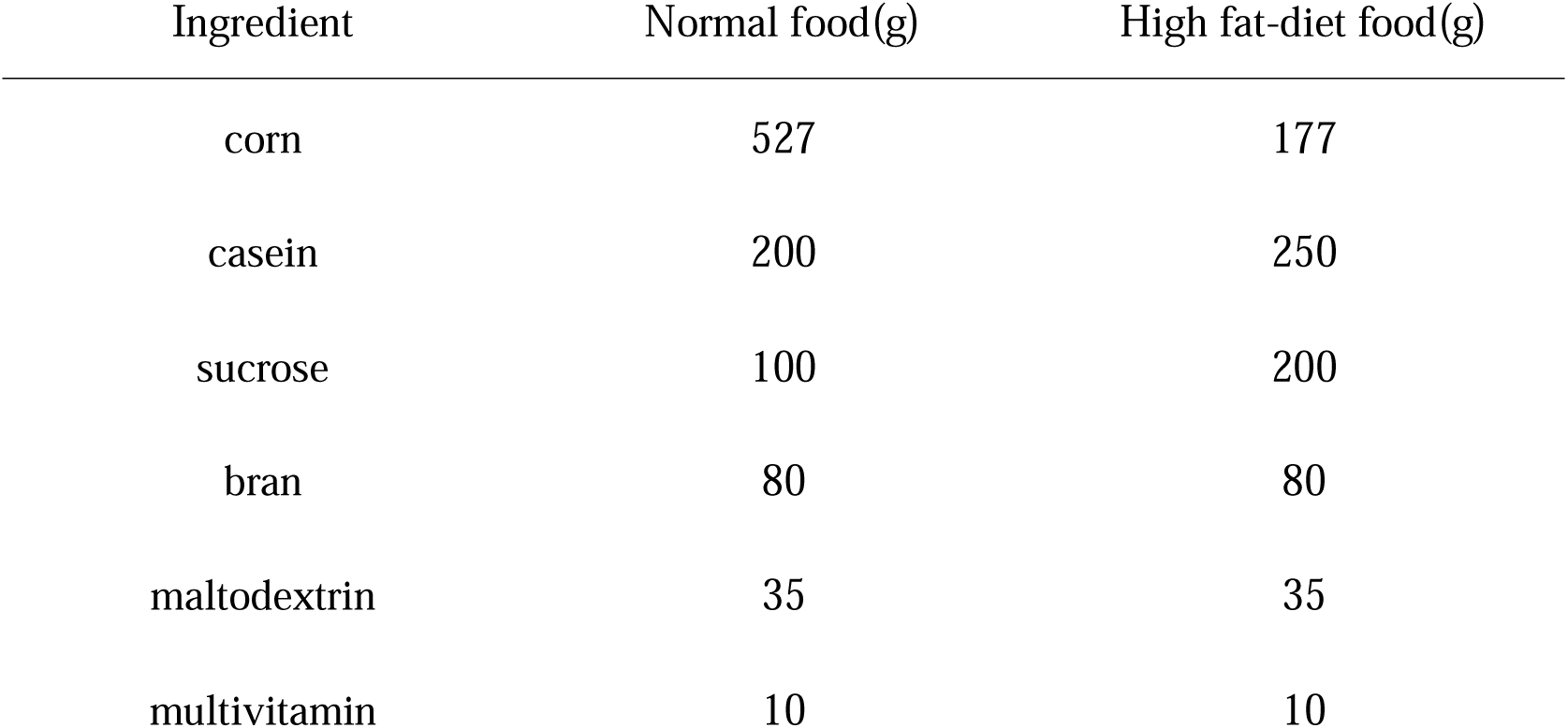
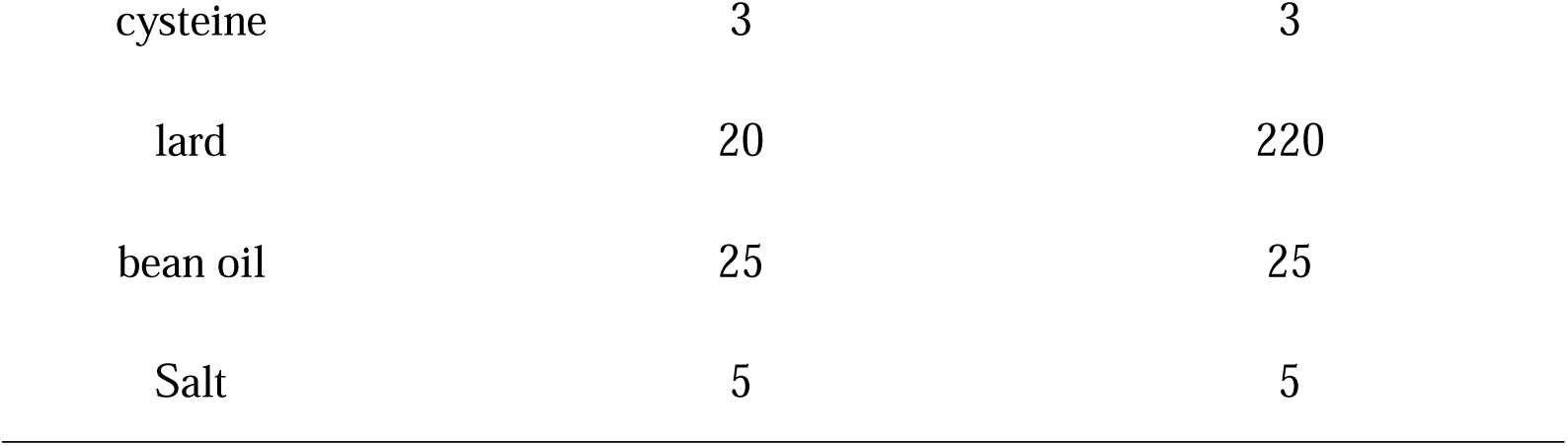
Food ingredient list per 1 kg.

### 2.2 Preparation of *Cyclocarya paliurus* polysaccharides

The leaves of C. *paliurus* were collected from Zhangjiajie, Hunan Province, China. The CCPP were extracted as described by Xie et al. with some modifications(29). The leaves of C. *paliurus* were air-dried and pulverized before extraction. First, 500 g dried C. *paliurus* leaves (60 mesh) were soaked with 2000 ml 80% ethanol (v/v) for 24 hours at room temperature, and the mixture was filtered. Second, the residues were dried in air and soaked with 4000 ml distilled water at 80[ twice for 150 min each time, then the mixture was filtered, and the filter liquor was retained. Third, the aqueous extract was concentrated to 20% of the original volume with a rotary evaporator at 60[ under vacuum. Fourth, 4 times volume of anhydrous ethanol was added to the concentrated liquid, the mixture was kept at 4[ for 24 hours, and the mixture was centrifugated at 10000 g for 15 min to remove the supernatant liquid. The above step was repeated 3 times. dilute *Cyclocarya paliurus* polysaccharides with the ratio of extraction: water =1:10, dilute and mix well, then store it at 4[.

### 2.3 *Cyclocarya paliurus* polysaccharides treatment

Mice (20±0. 5 g) were randomly grouped to normal diet (ND), high fat diet (HFD), and high fat diet plus *Cyclocarya paliurus* polysaccharides (CP) groups (n=6). ND feed normal diet, normal water, HFD feed high fat diet, normal water, CP feed high fat diet, add CCPP to drinking water. The concentration of C*. paliurus* leaves extract was 100mg/ml, and each mice received 400mg of plant extract through drinking water every day. Each group were weighed and fed every Sunday morning. After twelve weeks of C. *paliurus* administration, blood samples were collected by orbital blooding. visceral adipose tissue and subcutaneous adipose tissues were weighed and collected. Mice livers and Jejunum and caecum were collected. The liver was fixed with 4% paraformaldehyde for 12h for HE staining, and the rest of the liver was preserved at -80[. Jejunum and caecum were rinsed with PBS buffer solution, then frozen with liquid nitrogen and stored at -80[ for qRT-PCR.

### 2.4 Gut microbiota transplantation

Male mice aged 8 weeks were divided into control group (ND), high-fat diet mice fecal bacteria solution transplantation group (FMT-HF) and CCPP mice fecal bacteria solution transplantation group (FMT-CP) (n=9 per diet group). FMT-HFD group and FMT-CP group were added antibiotics (ampicillin 0.5g/L, neomycin 0.25g/L and metronidazole 0.5g/L) in drinking water before fecal bacteria solution transplantation, and antibiotics (Ampicillin 1g/L, neomycin 0.5g/L, vancomycin 0.5g/L and metronidazole 1g/L) 200μl by oral gavage for 4 days to clear gut microbiota(6). All antibiotics were purchased from Sigma-Aldrich (Shanghai) Trading Co.Ltd. and drinking water was prepared daily. After antibiotic treatment, 3 fecal samples were randomly collected in each cage of MT-HF and MT-CP groups, placed in a 1.5ml EP tube, and 1ml sterile PBS buffer was added to mix the fecal samples, and the mixture was thoroughly ground. The fecal bacteria solution was extracted by centrifugation at 3000r/min for 3 minutes. The bacterial content in feces was estimated by dilution coating plate method in order to tested the effect of antibiotic treatment.

8-week-old male donor mice (n=4 per diet group) were fed with HFD, or HFD add CCPP for 15weeks. After 4weeks of feeding, stools were collected daily for the subsequent 2 months under a laminar flow hood in sterile conditions. Stools from donor mice of each diet group were pooled and 200 mg was resuspended in 1 ml of PBS buffer solution, centrifuge the mixture at 3000r/min for 3 min. The supernatant was collected as transplant material, this process takes place in a biosafety cabinet. Fresh transplant material was prepared on the same day of transplantation within 15 min before oral gavage to prevent changes in bacterial composition. FMT-HFD and FMT-CP group were fed inoculated fresh transplant material (0.2ml for each mouse) three times a week by oral gavage for 2 months, before being killed for subsequent analysis(31).

### 2.5 Serum biochemical parameters

Serum samples were separated after centrifugation at 3000 rpm for 10 min under 4 °C. Epoch microplate spectra photometer was used to test serum biochemical parameters. Biochemical kits for cholesterol (CHOL), triglycerides (TG), low density lipoprotein (LDL), high density lipoprotein (HDL) and glucose (GLU)were purchased form Nanjing Jiancheng Bioengineering Institute (Nanjing, China).

### 2.6 Quantitative real-time reverse-transcription PCR

Total RNA from small intestine was isolated from liquid nitrogen frozen and ground tissues with TRIZOL regent (Sigma, USA). Equal amounts of total RNA were used to synthesize cDNA with the *PrimeScript*™ RT reagent Kit with gDNA Eraser (Takara RR047A, China). Quantitative real-time reverse-transcription PCR (qRT–PCR) was performed in triplicate using TB Green, 96-well plates and the CFX96 Real-Time PCR Detection System (bio-red). Each well was loaded with a total of 25μl containing 2μl of cDNA, 2μl of target primers, 8. 5μl of water and 12. 5μl of TB Green Premix Ex Taq II. β-actin was chosen as the house-keeping gene to normalize target gene levels. Hot-start PCR was performed for 50 cycles, with each cycle consisting of denaturation for 5 s at 95 [, annealing for 30 s at 60[ and elongation for 30 s at 60[. Relative quantification was done using the 2^-( ΔΔCT)^ method(32), where ΔΔ*Ct*= (*CtTarget* - *Ctactin*) treatment - (*CtTarget* - *Ct*β*-actin*) control. Relative expression was normalized and expressed as a ratio to the expression in the control group. The primers used are shown in Table 2.

**Table2:**
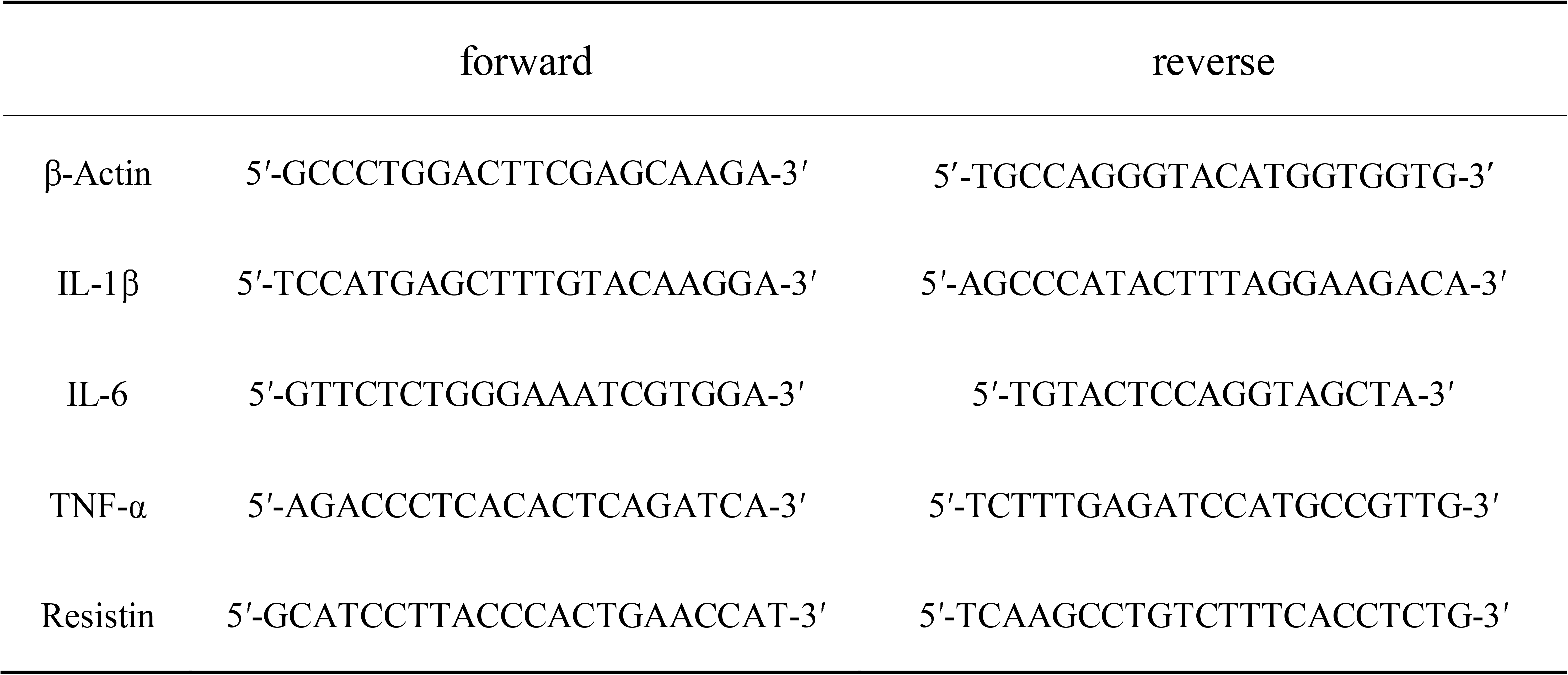
Gene primer sequence.

### 2.7 Hematoxylin and eosin (HE) staining

The liver tissue chunks preserved in 4% paraformaldehyde were dehydrated and transparent, dipped in wax and embedded in a dehydrator. After solidification, the slices were sliced by a slicer (thickness 4-6μm), and the slices were attached to the slides and baked at 65 [ for about 2 hours. HE staining was conducted according to standard method(33).

### 2.8 Mice gut microbes 16SrRNA genes high-throughput sequencing

After the sacrifice of each group of mice, cecal contents were collected, quick-frozen with liquid nitrogen, stored in dry ice and transported to GENEWIZ Biotechnology Co. Ltd. (Suzhou, China) for detection.

Total genome DNA from mice cecal contents was extracted according to manufacturer’s protocols. DNA concentration was monitored by *Equalbit* dsDNA HS Assay Kit. 20-30 ng DNA was used to generate amplicons. V3 and V4 hypervariable regions of prokaryotic 16S rDNA were selected for generating amplicons and following taxonomy analysis. GENEWIZ designed a panel of proprietary primers aimed at relatively conserved regions bordering the V3 and V4 hypervariable regions of bacteria and Archaea16S rDNA. Then, a linker with Index is added to the end of the PCR product of 16S rDNA by PCR for NGS sequencing, the library is purified with magnetic beads, and the concentration is detected by a microplate reader and the fragment size is detected by agarose gel electrophoresis. Detect the library concentration by a microplate reader. The library was quantified to 10nM, and PE250/FE300 paired-end sequencing was performed according to the Illumina MiSeq/Novaseq (Illumina, San Diego, CA, USA) instrument manual. The MiSeq Control Software (MCS)/Novaseq Control Software (NCS) Read sequence information.

Double-end sequencing of positive and negative reads the first of the two joining together to filter joining together the results contained in the sequence of N, retains the sequence length is larger than 200 bp sequence. After quality filter, purify chimeric sequences, the resulting sequence for OTU clustering, use VSEARCH clustering (1.9.6) sequence (sequence similarity is set to 97%), than the 16 s rRNA reference database is Silva, 132. Then use RDP classifier (Ribosomal Database Program) bayesian algorithm of OTU species taxonomy analysis representative sequences, and under different species classification level statistics community composition of each sample.

Based on OTU analysis results are obtained, using the method of random sampling sample sequences is flat, calculate Shannon, Chao1 alpha diversity index, community species Abundance. PCoA (principal co - ordinates analysis) based on the distance between the matrix Brary - Curtis. LEfSE analysis is more group, the differences between the species and the branching tree diagram shows the hierarchy of evolution between group differences of microbial community structure and species.

### 2.9 Statistical Analysis

Data are expressed as the mean ± sem. Differences date were assessed using the unpaired two-tailed Student’s t-test (Execl software). All statistical analyses were performed by using the one-way analysis of variance (ANOVA) to test homogeneity of variances via Levene’s test, A P value of < 0. 05 was considered significant.

## 3. Result

### 2.1 CCPP alleviates adipose accumulation in HF diet-fed mice

To determine the effect of CCPP on HF diet-fed mice, body weight, food intake, subcutaneous adipose tissue, and visceral adipose tissue were weighed (Figure1A-D). As expected, HFD significantly increased the body weight (P<0.001) (Figure1A), and food intake no significant difference (P>0.05) (Figure1B). Similarly, high fat diet significantly increased the weight of subcutaneous adipose tissue (P<0.001) and visceral adipose tissue (P<0.001). CCPP treatment tended to alleviate HF-induced weight gain, the difference was significant (P<0.05) (Figure1A). Notedly, the weight of subcutaneous adipose tissue and visceral adipose tissue was reduced in CP group compared to the HF group (P<0.05) (Figure1C, D). Images of representative ND, HFD or CP group mice subcutaneous adipose tissue and visceral adipose tissue during dissection at week 12 (Figure1C, D).

**Figure1.**
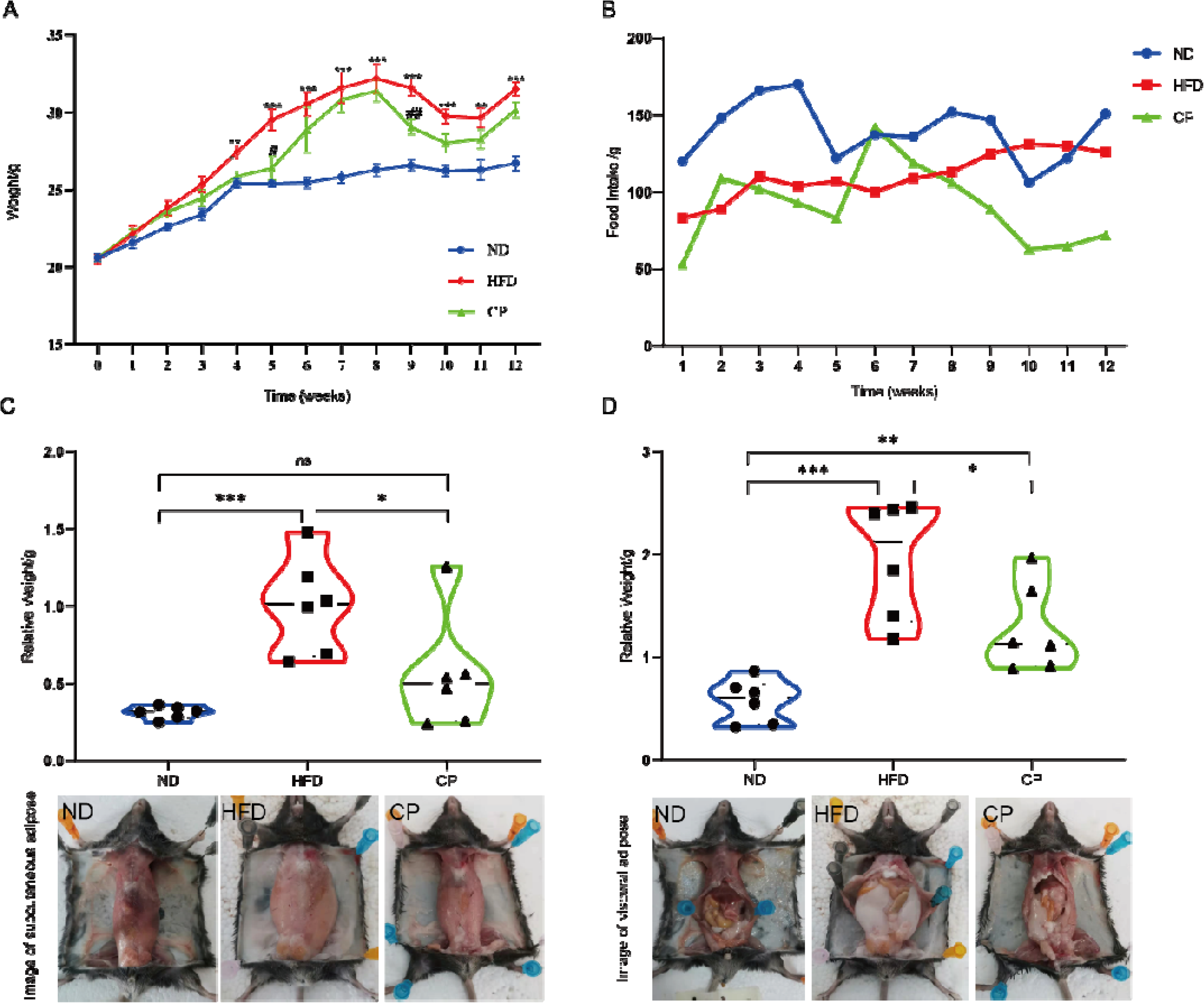
CCPP treatment improved lipid metabolism in high fat diet fed mice. Changes of body weight (A, ×±x) and average food intake (B) in each group of mice (n=6).(C)Mice subcutaneous adipose tissue weight and subcutaneous adipose tissue appearances of mice in the ND, HFD and CP groups. (D) Mice visceral adipose tissue weight and visceral adipose tissue appearances of mice in the ND, HFD and CP groups. Values are presented as mean ± SEM. Differences were assessed by student’s T test and denoted as follows: *P<0.05; **P<0. 01; ***P<0.001; ns P>0.05. * in the section of (A) body weight means the difference compared with the control group. #P<0.05; ##P<0. 01. #means the difference between CP and HFD group.

### 3.2 CCPP treatment improves hepatocyte steatosis and serum concentrations of lipid in HFD-fed mice

The incidence of steatohepatitis induced by chronic feeding with HFD was 100% in the HFD group. The hepatocytes had an abundant cytoplasm and were centrally located around the nuclei. The cords of the hepatocytes were arranged radially around the central vein. By contrast, the mice in the model group had disorganized cords, deformed and compressed hepatocytes, and cytoplasmic accumulation of lipid droplets with varied size, number and shape. In the CP group, liver damage was less severe than the HFD group. Most of the hepatocytes had a normal ultrastructure, intact cell membranes and less lipid droplet accumulation (Figure2A).

**Figure 2.**
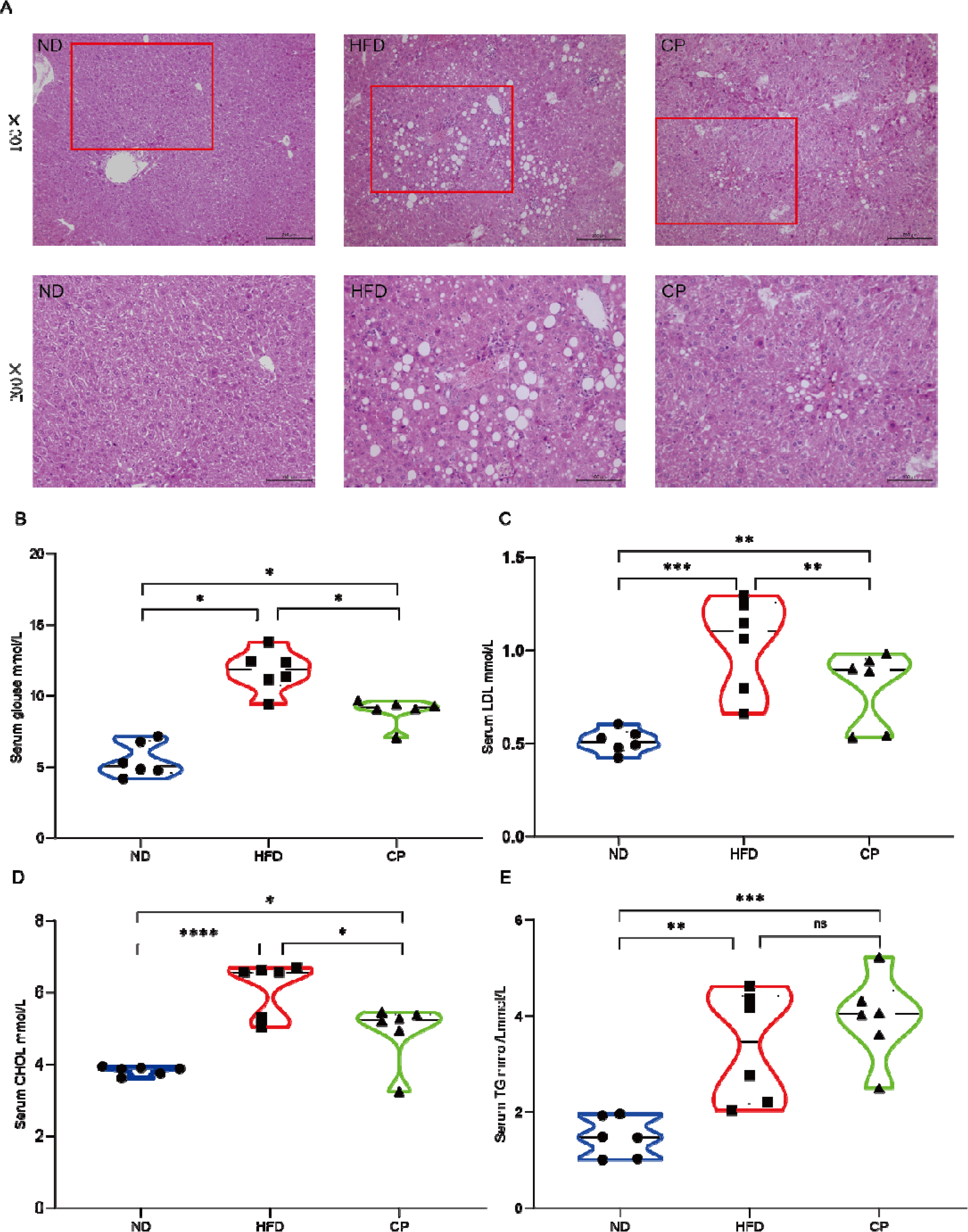
CP attenuates high-fat diet-induced liver injury. (A)Liver H. E staining (×200). (B-E) Serum concentrations of blood glucose, LDL, CHOL and TG in mice (n=6). Values are presented as mean ± SEM. Differences were assessed by student’s T test and denoted as follows: *P<0.05; **P<0. 01; ***P<0.001; ns P>0.05.

Next, we analyzed serum concentrations of lipid (Figure2B-E), and found that HF diet increased serum levels of GLU and TG (P<0.05), interestingly, CHOL and LDL levels increased significantly (P<0.001). CCPP treatment significantly reduced serum levels of CHOL, TG, LDL and GLU compared to the HFD group (P<0.05). HDL levels were also measured in this study, but there was no difference between ND and HFD. Collectively, CCPP treatment improved steatohepatitis, lipid accumulation, GLU, LDL, TC and TG metabolism, while it had little effect on HDL.

### 3.3 CCPP reduces inflammation in intestinal tract of HFD-fed mice

Gut microbiota disturbance induced by high fat diet increased the cell wall lysis component lipopolysaccharide (LPS) of Gram-negative bacteria, and LPS, as an inflammatory trigger factor, combined with TLR4 receptor to activate NF-⎢B signaling pathway and promote the synthesis and release of inflammatory factors TNF-α, IL-6 and IL-1β(34, 35). This can lead to a state of inflammation in the intestinal tissue. we measured expression of several genes related to inflammation in the gut tissue, including IL-1β, IL-6, TNF-α, MCP-1 and Resistin. IL -1β, IL -6 and TNF-α were significantly increased compared with the ND group (Figure 3). Meanwhile, CCPP reduced mRNA abundance of IL-6 compared to the HFD group (P<0.05). Also, CCPP significantly (P<0.001) downregulated mRNA expression of IL-1β and TNF-α in the liver.

**Figure 3.**
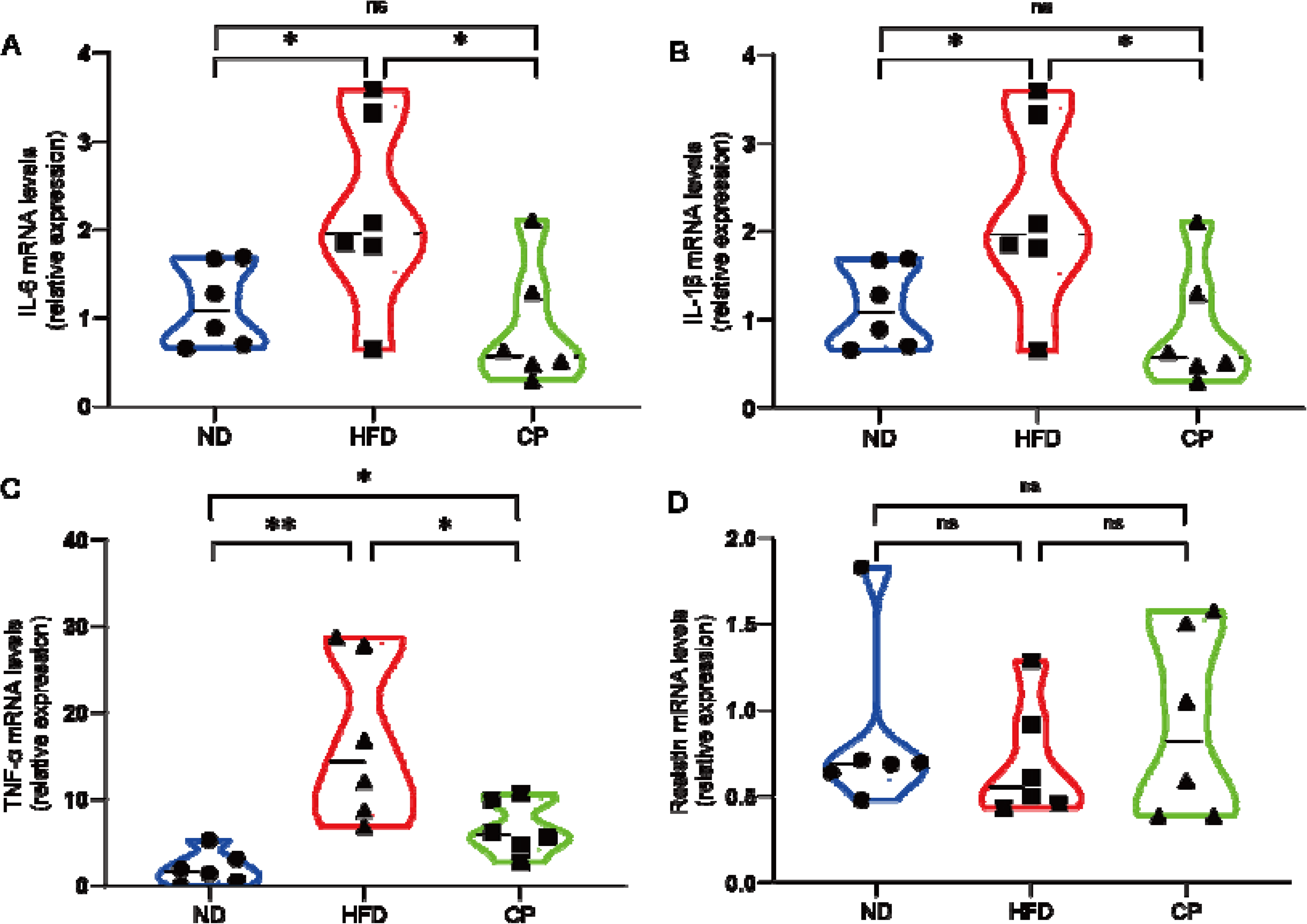
CP treatment reversed gene expressions of Inflammatory factor in the gut of HF diet-fed mice. IL-1β, IL-6, TNFα and Resistin mRNA abundances were determined by real-time PCR analysis and relative gene pressions were normalized with β-actin (n=6). Values are presented as mean ± SEM. Differences were assessed by student’s T test and denoted as follows: *P<0.05; **P<0. 01; ***P<0.001; ns P>0.05.

### 3.4 CCPP ameliorates gut microbiota disorder induced by high fat diet-fed mice

Gut microbiota is highly associated with obesity and lipid metabolism, so we further determined the composition of fecal microbiota by sequencing intestinal microbiota 16S rRNA. In this study, an average of 74, 742 raw reads were generated from each sample. After removing the low-quality sequences, 70, 503 clean tags were clustered into OTUs. β diversity analysis can reflect whether there is difference between different samples. UniFrac distance-based principal coordinate analysis (PCoA) revealed distinct clustering of intestinal microbe communities for each experimental group. Remarkable changes in the microbiota community structure were induced by both HFD and CCPP intervention. As shown in Figure 4A, the abscissa is PC1 with a contribution rate of 30.73%, and the ordinate is PC2 with a contribution rate of 18.29%. The microorganisms in ND group and HFD group were far apart, and there was a significant distance between CP group and HFD group, which is an indication that CCPP consumption induced similar microbial composition changes.

**Figure 4.**
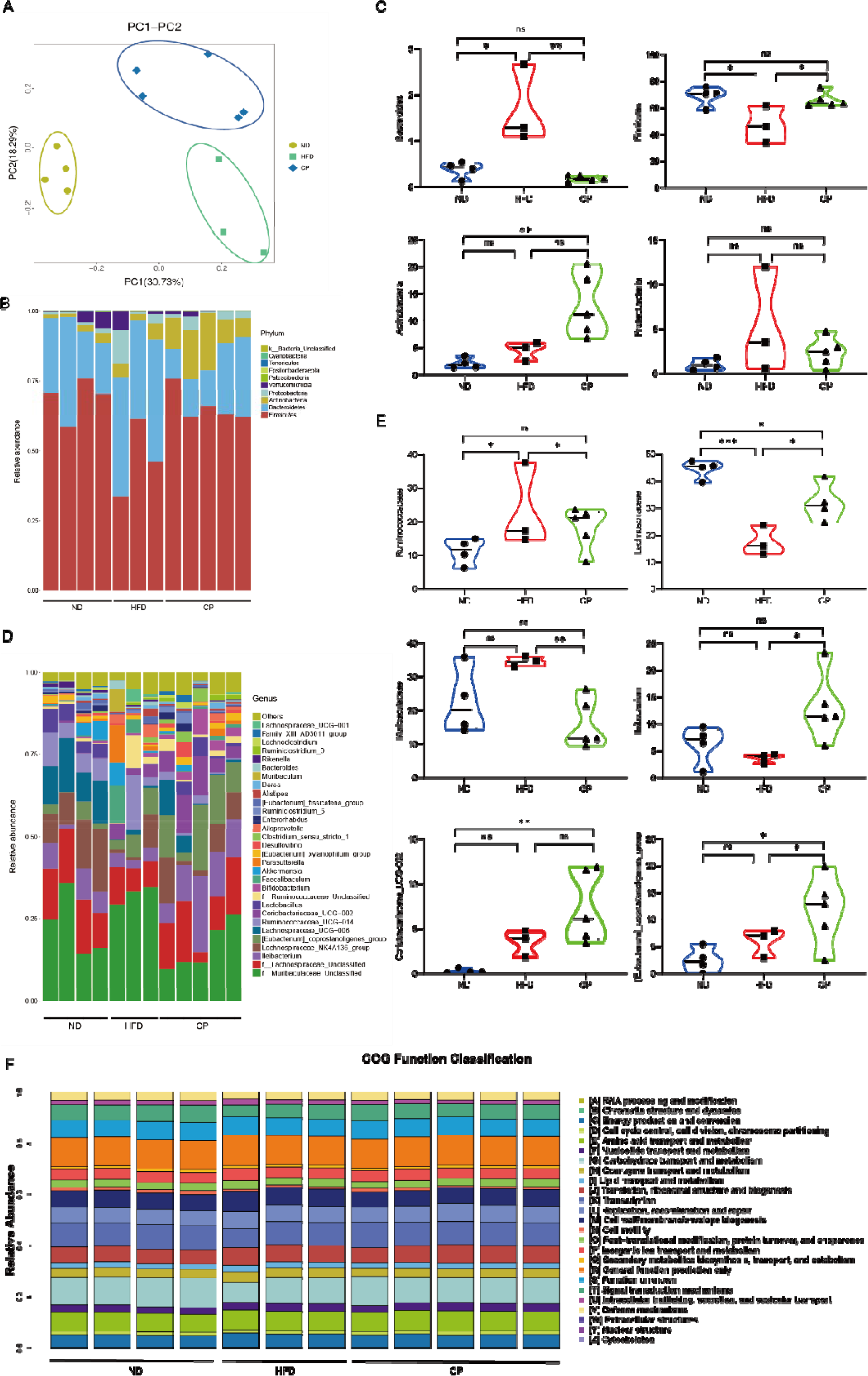
CP alters microbiota composition and microbiota function in HFD-fed mice. (A) Scatter plot of the principal coordinate analysis (PCoA) score showing the similarity of the 12 bacterial communities based on the Unifrac distance. Principal components (PCs) 1 and 2 explained 30.73% and 18.29% of the variance, respectively; (B-C) Relative abundance of species at the phylum level. (D-E) Relative abundance of species at the genus level.

Through comparative analysis of the sequence of the sequenced samples, we studied the fecal microbial community structure of CP-treatment mice, and compared with ND and HFD groups. As shown in Figure 4B and 4C, the intestinal flora of the three samples at phylum level is mainly *Firmicutes*, *Bacteroidetes*, *Actinobacteria* and *Proteobacteria*. It is worth noting that the composition of mice gut microbiota at phylum level was different from that of other studies, especially the ratio of *Firmicutes* to *Bacteroidetes*, in which *Firmicutes* abundance increased in ND group and CP group, while *Firmicutes* abundance decreased in HFD group. On the contrary, *Bacteroidetes* abundance increased in HFD group and decreased in CP group and ND group. The relative abundance of *Firmicutes* in ND group, HFD group and CP group was 68.93%, 47.19% and 66.02%, respectively. Compared with HFD group, the relative abundance of *Firmicutes* in ND group was significantly different (P<0.05). The relative abundance of *Bacteroidetes* was 25.20% in ND group, 17.99% in CP group and 40.23% in HFD group, 15.03% lower than that of ND group. We found that the relative abundance value of actinomycetes increased in the CCPP feeding group (12.94%), which was significantly different from the ND group (P=0.0095).

The relative abundance of species at the genus level is shown in Figure 4D and 4E. At the genus level, *Muribaculaceae_Unclassified*, *Lachnospiraceae_Unclassified*, *Ileibacterium*, *Lachnospiraceae_NK4A136_group* are the most important four genera. Among them, *Lachnospiraceae_Unclassified* is the second dominant bacterium. Many studies have found that the bacteria belonging to *Lachnospiraceae* is tightly associated with the production of SCFAs(30, 31, 36). The relative abundance of *Lachnospiraceae_Unclassified* in ND group, HFD group and CP group was 14.96%, 8.81% and 12.70%, respectively. Compared with normal control group, the relative abundance of *Lachnospiraceae_Unclassified* in HFD group was significantly different (P<0.05), and CP feeding could inhibit the decrease of the relative abundance of *Lachnospiraceae_Unclassified*. Consistently, the CCPP-treatment mice were characterized by a higher abundance of *Ileibacterium* and *[Eubacterium]_coprostanoligenes_group*, of which Illi was reported to be beneficial(37).

COG pathways were further annotated according to the microbiota compositions by PICRUSt analysis, and the results showed that microbiota-mediated genes were mainly involved in Energy production and conversion (20,526,768genes), Amino acid transport and metabolism (30,885,519genes), Carbohydrate transport and metabolism (42,773,412genes), Translation, ribosomal structure and biogenesis (25,750,116genes), Lipid transport and metabolism (9,095,565genes) and Coenzyme transport and metabolism (14,742,277genes) (Figure 4F). Among them, HF diet markedly reduced Amino acid transport and metabolism, Nucleotide transport and metabolism, Lipid transport and metabolism and Cell motility. CCPP treatment reprogramed the Carbohydrate transport and metabolism, Translation, ribosomal structure and biogenesis and repair.

### 3.5 Microbiota transplantation from CP-treated mice fails to affect lipid metabolism in HF diet-fed mice

We then sought to investigate whether microbiota transplantation from CP-treated mice could alleviate lipid accumulation in HF diet-fed mice. High fat diet-fed mice received regular or CP extracts water for 4 weeks (n=4), respectively. Then fecal samples were collected and transplanted into antibiotics-treated mice (n=9). Compared with the control group of mice without antibiotic treatment, the number of bacteria in FMT-HFD group and FMT-CP group decreased significantly, indicating that antibiotics kill most intestinal flora.

Microbiota-transplanted mice were further fed high fat diet for 8 weeks. As shown in Figure5A. The average body weight of FMT-CP group was lower than that of FMT-HFD group, but the difference was not statistically significant(P>0.05). Food intake no significant difference (Figure5B). At week 8 of microbiota transplantation, we sacrificed the mice. Subcutaneous adipose tissue and visceral adipose tissue quality of mice in each group is shown in Figure5C-F, Compared with FMT-HFD, the weight of subcutaneous adipose tissue in FMT-CP was significantly reduced(P<0.05), but there was no difference in visceral adipose tissue (P>0.05). Interestingly, microbiota transplantation from CP-treated mice decreased serum levels of GLU (P<0.05) and TG (P<0.01), but had little effect on serum levels of TC, LDL, and CHOL (P>0.05). These results indicated that microbiota transplantation from CP-treated mice slightly improves lipid metabolism in HFD mice.

**Figure5.**
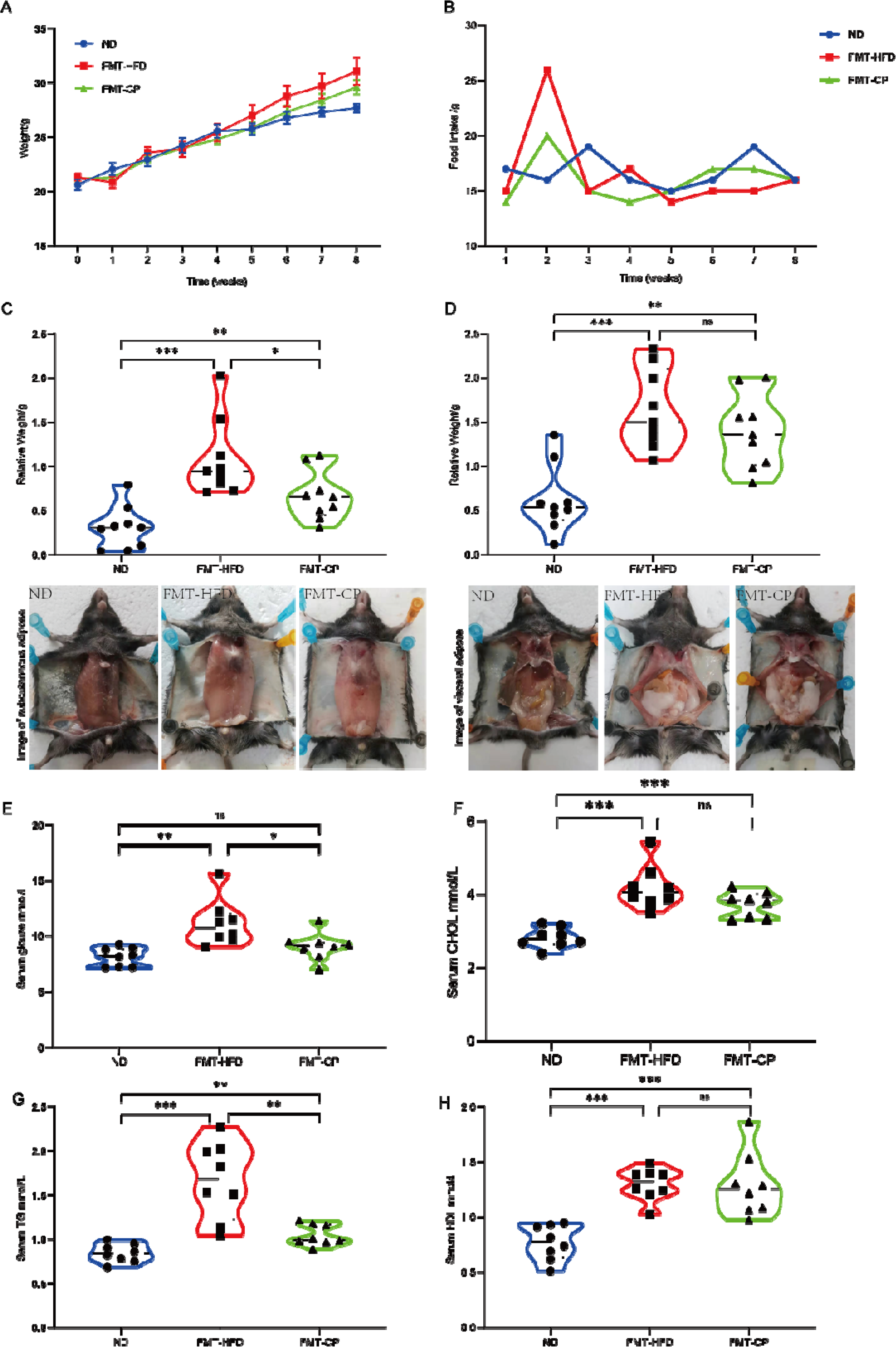
Microbiota transplantation from CCPP-treated mice improve lipid metabolism in HF-diet fed mice. Changes of body weight (A, ×±x) and average food intake (B) in each group of mice (n=9). (C)Subcutaneous adipose tissue appearances of mice in the ND, HFD and CP groups. (D)Mice subcutaneous adipose tissue weight. (E)visceral adipose tissue appearances of mice in the ND, HFD and CP groups. (F)Mice visceral adipose tissue weight. (G-J) Serum concentrations of blood glucose, CHOL, LDL, TG and HDL in mice (n=6). Values are presented as mean ± SEM. Differences were assessed by student’s T test and denoted as follows: *P<0.05; **P<0. 01; ***P<0.001; ns P>0.05.

**Figure6:**
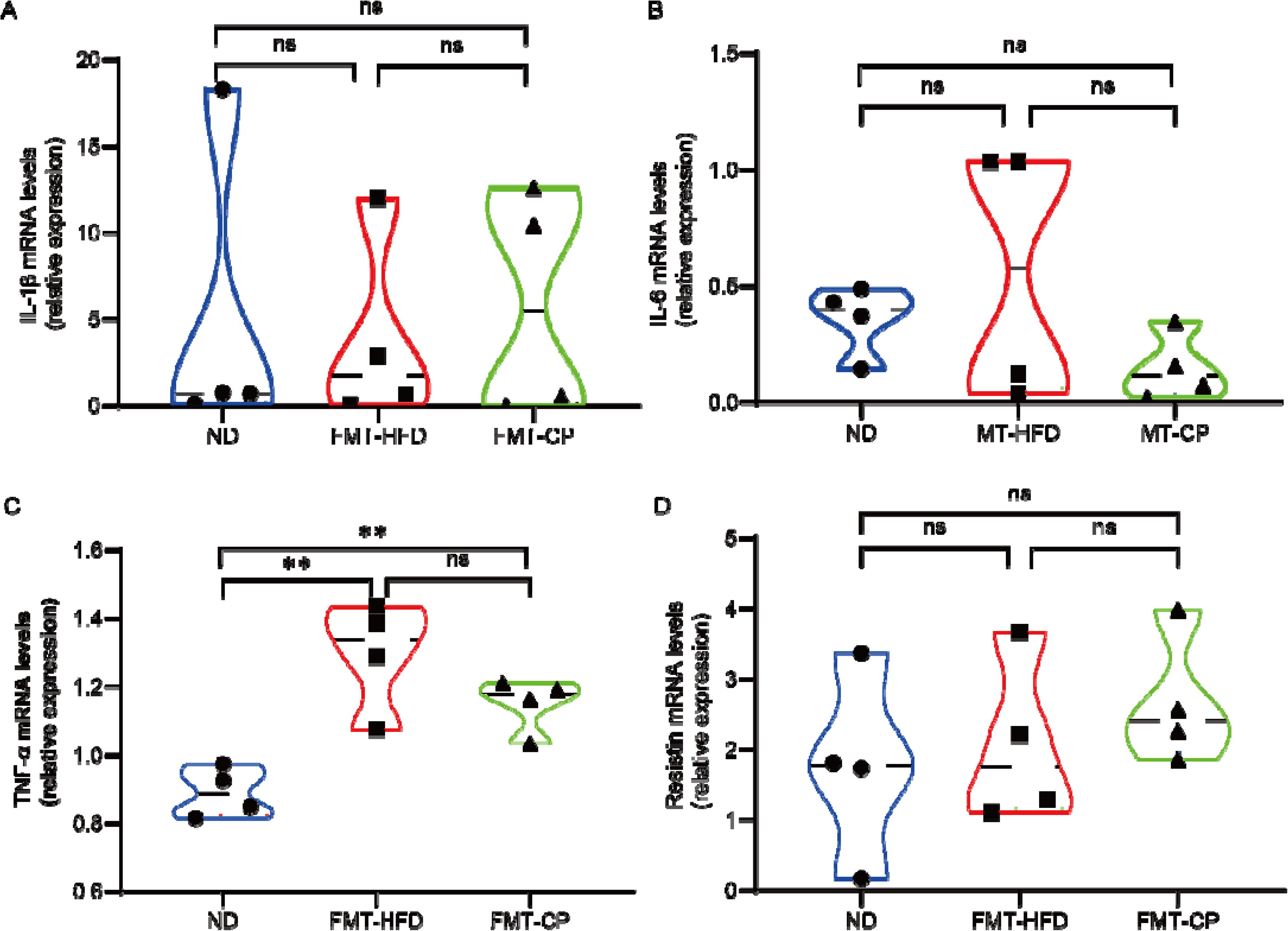
Microbiota transplantation treatment reversed gene expressions of Inflammatory factor. IL-1β, IL-6, TNFα and Resistin mRNA abundances were determined by real-time PCR analysis and relative gene pressions were normalized with β-actin (n=4). Values are presented as mean ± SEM. Differences were assessed by student’s T test and denoted as follows: *P<0.05; **P<0. 01; ***P<0.001; ns P>0.05.

### 3.6 Microbiota transplantation from CP-treated mice fail to reduces inflammation in intestinal tract of HF diet-fed mice

In order to explore whether microbiota transplantation can reduce the expression of intestinal inflammatory factors, we detected the mRNA expression of IL-1β, IL-6, TNF-α and Resistin. Interesting, compared with ND group, microbiota transplantation could reduce the mRNA expression of IL-1, IL-6 and Resistin in FMT-CP and FMT-HFD groups(P>0.05). However, microbiota transplantation can’t reduce TNF-α mRNA expression, compare with ND group, FMT-CP and FMT-HFD group significantly improve TNF-α mRNA expression(P<0.01).

## 4. Discussion

We and others have previously shown that gut microbiota are sensitive to CCPP exposure and CCPP treatment markedly shapes or reverses gut microbiota in healthy or obese animal models(23, 30, 38, 39). In these studies, however, no direct evidence was obtained to support that gut microbiota mediate the CCPP-induced beneficial potentials. Thus, we propose whether the anti-obesity functions of CCPP depend on the gut microbiota. Our results showed that CCPP has a protective effect against HFD-induced obesity, consistent with previous studies(23, 29, 30). CCPP supplementation restored the gut microbiota composition and function which was destroyed by the HFD. Moreover, the anti-obesity and microbiota-modulating effects of CCPP are transferrable through FMT.

We found a significant decrease of inflammation-related cytokines IL-1β, IL-6, TNF-α and Resistin in the CP treatment group. Thus, CCPP treatment significantly reverses systemic GLGI and inflammatory factors expression in HF-fed mice. This was in accordance with previous study(38).

Accumulating evidence has revealed that gut microbiota was closely related to the health(40, 41). Long-term dietary changes will lead to profound alterations in gut microbiota composition. Many dietary supplements have been used to prevent or alleviate metabolic disease by modulating the gut microbiota. Here, we found that the composition of the gut microbiota was notably altered by CCPP. *Firmicutes*, *Bacteroidetes*, *Actinobacteria* and *Proteobacteria* were the main phyla in all groups, but *Bacteroidetes* and *Proteobacteria* were more abundant in the HFD group, and *Firmicutes* and *Actinobacteria* were more abundant in the CCPP group. CCPP could promote *Firmicutes* growth and decreased proportion of *Bacteroidetes* in CCPP-treated mice, this is in agreement with previous reports(30). Intriguingly, previous study reported that Ganoderma lucidum polysaccharide reduced *Firmicutes*/*Bacteroidetes* ratio in HFD-fed obesity mice(42). It was recently found that a low *Bacteroidetes*-to-*Firmicutes* ratio was correlated with body weight loss(43). At the family level, several beneficial genera responding to CCPP treatment were identified, especially the *Lachnospiraceae*. *Lachnospiraceae* is one of the most abundant families from the order Clostridiales in gut environment, and had been associated with the maintenance of gut health. *Lachnospiraceae* species hydrolyze starch and other sugars to produce butyrate and other SCFAs(44, 45), and some studies showed that *Lachnospiraceae* may also be specifically associated with type 2 diabetes (T2D) in both humans and mouse models(46, 47), however a metagenome-wide association study revealed a loss of several butyrate-producing bacteria in faecal samples from T2D patients(48), suggesting a potential protective role uniquely for butyrate. At the genus level, *Coriobacteriaceae_UCG-002* was abundant in the CCPP group. *Coriobacteriaceae_UCG-002*, although belonging to the *Actinobacteria* phylum, has been reported to be beneficial for improved glucose tolerance and insulin sensitivity(49). In the present study, CCPP treatment restored gut microbiota dysbiosis by preventing the loss of *Firmicutes*, *Lachnospiraceae* and increasing *Coriobacteriaceae_UCG-002* to bring it closer to ND group. We did not measure the levels of SCFAs in either feces or the serum due to the limited number of samples collected, which we recognize as a limitation in our study.

Concomitant with the gut microbial structure alteration, we observed an improvement in bacterial gene functions after CCPP supplementation. Among them, HF diet markedly reduced Amino acid transport and metabolism, Nucleotide transport and metabolism, Lipid transport and metabolism and Cell motility. After CCPP treatment the Carbohydrate transport and metabolism, Translation, ribosomal structure and biogenesis and repair pathways were increasing. These findings are consistent with increased bacterial production of SCAFs and may contribute to the treatment of obesity.

A key question is whether the important anti-obesity effects of CCPP on mice are dependent on the alteration of the gut microbiota. To test this question, we performed FMT to investigate the causal role of microbiota. We demonstrated that transplantation of CCPP-treated mouse feces to HFD-fed recipients, improved their metabolic. Microbiota transplantation from CP-treated mice could alleviate subcutaneous adipose tissue weight, GLU and TG levels in HF diet-fed mice. In addition, our further study found that the effect of CCPP on reducing the expression of inflammatory factors could be partially transferred by gut microbiota transplantation. The expression of inflammatory factor IL-1 in FMT-CP group was significantly lower than that in FMT-HFD group. However, there was no difference in the expression of other inflammatory factors such as IL-6 and TNF-α.

## 6. Acknowledgement

We would like to thank all the members of Chen’s laboratory for their help and useful discussions, as well as Professor Chen and Professor Liu for their helpful guidance. This work was supported by the Research Fund of General project of Basic Research Plan of Shanxi Province in 2021(Project Number: 20210302123239) and Shanxi College Student Training Program for Innovation and Entrepreneurship (S202117114001).

## References

1. Organization WH. Obesity and Overweight 2020 [Available from: https://www.who.int/news-room/fact-sheets/detail/obesity-and-overweight.

2. Organization WH. Noncommunicable diseases 2017 [Available from: http://www.who.int/mediacentre/factsheets/fs355/en/.

3. Grundy SM. Obesity, metabolic syndrome, and cardiovascular disease. (0021-972X (Print)).

4. Ghaben AL, Scherer PA-O. Adipogenesis and metabolic health. (1471-0080 (Electronic)).

5. Ticinesi A, Lauretani F, Milani C, Nouvenne AA-O, Tana C, Del Rio DA-O, et al. Aging Gut Microbiota at the Cross-Road between Nutrition, Physical Frailty, and Sarcopenia: Is There a Gut-Muscle Axis? LID - 10.3390/nu9121303 [doi] LID - 1303. (2072-6643 (Electronic)).

6. Yin J, Li Y, Han H, Chen S, Gao J, Liu G, et al. Melatonin reprogramming of gut microbiota improves lipid dysmetabolism in high-fat diet-fed mice. (1600-079X (Electronic)).

7. Dalby MJ, Ross AW, Walker AW, Morgan PJ. Dietary Uncoupling of Gut Microbiota and Energy Harvesting from Obesity and Glucose Tolerance in Mice. (2211-1247 (Electronic)).

8. Cani PD, Neyrinck Am Fau - Fava F, Fava F Fau - Knauf C, Knauf C Fau - Burcelin RG, Burcelin Rg Fau - Tuohy KM, Tuohy Km Fau - Gibson GR, et al. Selective increases of bifidobacteria in gut microflora improve high-fat-diet-induced diabetes in mice through a mechanism associated with endotoxaemia. (0012-186X (Print)).

9. Chevalier C, Stojanović O, Colin DJ, Suarez-Zamorano N, Tarallo V, Veyrat-Durebex C, et al. Gut Microbiota Orchestrates Energy Homeostasis during Cold. (1097-4172 (Electronic)).

10. Fabbiano S, Suárez-Zamorano N, Chevalier C, Lazarević V, Kieser S, Rigo D, et al. Functional Gut Microbiota Remodeling Contributes to the Caloric Restriction-Induced Metabolic Improvements. (1932-7420 (Electronic)).

11. Gomes AC, Hoffmann C, Mota JF. The human gut microbiota: Metabolism and perspective in obesity. (1949-0984 (Electronic)).

12. Ridaura VK, Faith Jj Fau - Rey FE, Rey Fe Fau - Cheng J, Cheng J Fau - Duncan AE, Duncan Ae Fau - Kau AL, Kau Al Fau - Griffin NW, et al. Gut microbiota from twins discordant for obesity modulate metabolism in mice. (1095-9203 (Electronic)).

13. Turnbaugh PJ, Ley Re Fau - Mahowald MA, Mahowald Ma Fau - Magrini V, Magrini V Fau - Mardis ER, Mardis Er Fau - Gordon JI, Gordon JI. An obesity-associated gut microbiome with increased capacity for energy harvest. (1476-4687 (Electronic)).

14. Goodman AL, Kallstrom G Fau - Faith JJ, Faith Jj Fau - Reyes A, Reyes A Fau - Moore A, Moore A Fau - Dantas G, Dantas G Fau - Gordon JI, et al. Extensive personal human gut microbiota culture collections characterized and manipulated in gnotobiotic mice. (1091-6490 (Electronic)).

15. Chang CJ, Lin CS, Lu CC, Martel J, Ko YF, Ojcius DM, et al. Ganoderma lucidum reduces obesity in mice by modulating the composition of the gut microbiota. (2041-1723 (Electronic)).

16. Huang F, Zheng X, Ma X, Jiang RA-O, Zhou W, Zhou S, et al. Theabrownin from Pu-erh tea attenuates hypercholesterolemia via modulation of gut microbiota and bile acid metabolism. (2041-1723 (Electronic)).

17. Li X, Watanabe K, Kimura I. Gut Microbiota Dysbiosis Drives and Implies Novel Therapeutic Strategies for Diabetes Mellitus and Related Metabolic Diseases. (1664-3224 (Print)).

18. Ruiz-Núñez B, Pruimboom L Fau - Dijck-Brouwer DAJ, Dijck-Brouwer Da Fau - Muskiet FAJ, Muskiet FA. Lifestyle and nutritional imbalances associated with Western diseases: causes and consequences of chronic systemic low-grade inflammation in an evolutionary context. (1873-4847 (Electronic)).

19. Hotamisligil GS. Inflammation and metabolic disorders. (1476-4687 (Electronic)).

20. Ewaschuk J, Endersby R Fau - Thiel D, Thiel D Fau - Diaz H, Diaz H Fau - Backer J, Backer J Fau - Ma M, Ma M Fau - Churchill T, et al. Probiotic bacteria prevent hepatic damage and maintain colonic barrier function in a mouse model of sepsis. (0270-9139 (Print)).

21. Li Q, Hu J, Xie J, Nie S, Xie MY. Isolation, structure, and bioactivities of polysaccharides from Cyclocarya paliurus (Batal.) Iljinskaja. (1749-6632 (Electronic)).

22. Liu W, Wu Y, Hu Y, Qin S, Guo X, Wang M, et al. Effects of Cyclocarya paliurus Aqueous and Ethanol Extracts on Glucolipid Metabolism and the Underlying Mechanisms: A Meta-Analysis and Systematic Review. (2296-861X (Print)).

23. Wang Q, Jiang C Fau - Fang S, Fang S Fau - Wang J, Wang J Fau - Ji Y, Ji Y Fau - Shang X, Shang X Fau - Ni Y, et al. Antihyperglycemic, antihyperlipidemic and antioxidant effects of ethanol and aqueous extracts of Cyclocarya paliurus leaves in type 2 diabetic rats. (1872-7573 (Electronic)).

24. Xie JH, Dong CJ, Nie SP, Li F, Wang ZJ, Shen MY, et al. Extraction, chemical composition and antioxidant activity of flavonoids from Cyclocarya paliurus (Batal.) Iljinskaja leaves. (1873-7072 (Electronic)).

25. Xie JH, Shen My Fau - Nie S-P, Nie Sp Fau - Liu X, Liu X Fau - Zhang H, Zhang H Fau - Xie M-Y, Xie MY. Analysis of monosaccharide composition of Cyclocarya paliurus polysaccharide with anion exchange chromatography. (1879-1344 (Electronic)).

26. Yang Z, Wang J, Li J, Xiong L, Chen H, Liu X, et al. Antihyperlipidemic and hepatoprotective activities of polysaccharide fraction from Cyclocarya paliurus in high-fat emulsion-induced hyperlipidaemic mice. (1879-1344 (Electronic)).

27. Xiong L, Ouyang KH, Jiang Y, Yang ZW, Hu WB, Chen H, et al. Chemical composition of Cyclocarya paliurus polysaccharide and inflammatory effects in lipopolysaccharide-stimulated RAW264.7 macrophage. (1879-0003 (Electronic)).

28. Liu X, Xie J, Jia S, Huang L, Wang Z, Li C, et al. Immunomodulatory effects of an acetylated Cyclocarya paliurus polysaccharide on murine macrophages RAW264.7. (1879-0003 (Electronic)).

29. Xie JH, Liu X Fau - Shen M-Y, Shen My Fau - Nie S-P, Nie Sp Fau - Zhang H, Zhang H Fau - Li C, Li C Fau - Gong D-M, et al. Purification, physicochemical characterisation and anticancer activity of a polysaccharide from Cyclocarya paliurus leaves. (1873-7072 (Electronic)).

30. Yao Y, Yan L, Chen H, Wu N, Wang W, Wang D. Cyclocarya paliurus polysaccharides alleviate type 2 diabetic symptoms by modulating gut microbiota and short-chain fatty acids. (1618-095X (Electronic)).

31. Borody TJ, Paramsothy S Fau - Agrawal G, Agrawal G. Fecal microbiota transplantation: indications, methods, evidence, and future directions. (1534-312X (Electronic)).

32. Livak KJ, Schmittgen TD. Analysis of relative gene expression data using real-time quantitative PCR and the 2(-Delta Delta C(T)) Method. (1046-2023 (Print)).

33. Cardiff RD, Miller CH, Munn RJ. Manual hematoxylin and eosin staining of mouse tissue sections. (1559-6095 (Electronic)).

34. Li H, Lelliott C Fau - Håkansson P, Håkansson P Fau - Ploj K, Ploj K Fau - Tuneld A, Tuneld A Fau - Verolin-Johansson M, Verolin-Johansson M Fau - Benthem L, et al. Intestinal, adipose, and liver inflammation in diet-induced obese mice. (1532-8600 (Electronic)).

35. Cani PD, Bibiloni R Fau - Knauf C, Knauf C Fau - Waget A, Waget A Fau - Neyrinck AM, Neyrinck Am Fau - Delzenne NM, Delzenne Nm Fau - Burcelin R, et al. Changes in gut microbiota control metabolic endotoxemia-induced inflammation in high-fat diet-induced obesity and diabetes in mice. (1939-327X (Electronic)).

36. Liu Z, Ai CA-O, Lin X, Guo X, Song SA-O, Zhu BA-O. Sea cucumber sulfated polysaccharides and Lactobacillus gasseri synergistically ameliorate the overweight induced by altered gut microbiota in mice. (2042-650X (Electronic)).

37. Wang Y, Ablimit N, Zhang Y, Li J, Wang X, Liu J, et al. Novel β-mannanase/GLP-1 fusion peptide high effectively ameliorates obesity in a mouse model by modifying balance of gut microbiota. (1879-0003 (Electronic)).

38. Li Q, Hu J, Nie Q, Chang X, Fang Q, Xie J, et al. Hypoglycemic mechanism of polysaccharide from Cyclocarya paliurus leaves in type 2 diabetic rats by gut microbiota and host metabolism alteration. (1869-1889 (Electronic)).

39. Wang X, Tang L, Ping W, Su Q, Ouyang S, Su JA-O. Progress in Research on the Alleviation of Glucose Metabolism Disorders in Type 2 Diabetes Using Cyclocarya paliurus. LID - 10.3390/nu14153169 [doi] LID - 3169. (2072-6643 (Electronic)).

40. John GK, Mullin GE. The Gut Microbiome and Obesity. (1534-6269 (Electronic)).

41. Gagen EJ, Padmanabha J, Denman SE, McSweeney CS. Hydrogenotrophic culture enrichment reveals rumen Lachnospiraceae and Ruminococcaceae acetogens and hydrogen-responsive Bacteroidetes from pasture-fed cattle. LID - fnv104 [pii] LID - 10.1093/femsle/fnv104 [doi]. (1574-6968 (Electronic)).

42. Xiao-Fang Z, Xiao-Qun D, Xi L. The effect of Cyclocarya Paliurus Polysaccharide(CP) on blood glucose and histomorphology of pancreas in diabetic mice. Acta Medicinae Sinica. 2010.

43. Amato KR, Yeoman Cj Fau - Kent A, Kent A Fau - Righini N, Righini N Fau - Carbonero F, Carbonero F Fau - Estrada A, Estrada A Fau - Gaskins HR, et al. Habitat degradation impacts black howler monkey (Alouatta pigra) gastrointestinal microbiomes. (1751-7370 (Electronic)).

44. Zhang J, Song L, Wang Y, Liu C, Zhang L, Zhu S, et al. Beneficial effect of butyrate-producing Lachnospiraceae on stress-induced visceral hypersensitivity in rats. (1440-1746 (Electronic)).

45. Vacca MA-OX, Celano GA-O, Calabrese FA-O, Portincasa PA-O, Gobbetti M, De Angelis MA-OX. The Controversial Role of Human Gut Lachnospiraceae. LID - 10.3390/microorganisms8040573 [doi] LID - 573. (2076-2607 (Print)).

46. Qin J, Li Y Fau - Cai Z, Cai Z Fau - Li S, Li S Fau - Zhu J, Zhu J Fau - Zhang F, Zhang F Fau - Liang S, et al. A metagenome-wide association study of gut microbiota in type 2 diabetes. (1476-4687 (Electronic)).

47. Kameyama K, Itoh K. Intestinal colonization by a Lachnospiraceae bacterium contributes to the development of diabetes in obese mice. (1347-4405 (Electronic)).

48. Qin J, Li R Fau - Raes J, Raes J Fau - Arumugam M, Arumugam M Fau - Burgdorf KS, Burgdorf Ks Fau - Manichanh C, Manichanh C Fau - Nielsen T, et al. A human gut microbial gene catalogue established by metagenomic sequencing. (1476-4687 (Electronic)).

49. Liu H, Zhang H, Wang X, Yu X, Hu C, Zhang X. The family Coriobacteriaceae is a potential contributor to the beneficial effects of Roux-en-Y gastric bypass on type 2 diabetes. (1878-7533 (Electronic)).

