## Supplementary figures and images for "*Cyclocarya paliurus* polysaccharides improve high fat diet-fed mice lipid dysmetabolism through gut microbiota"

### supplementary figure 1

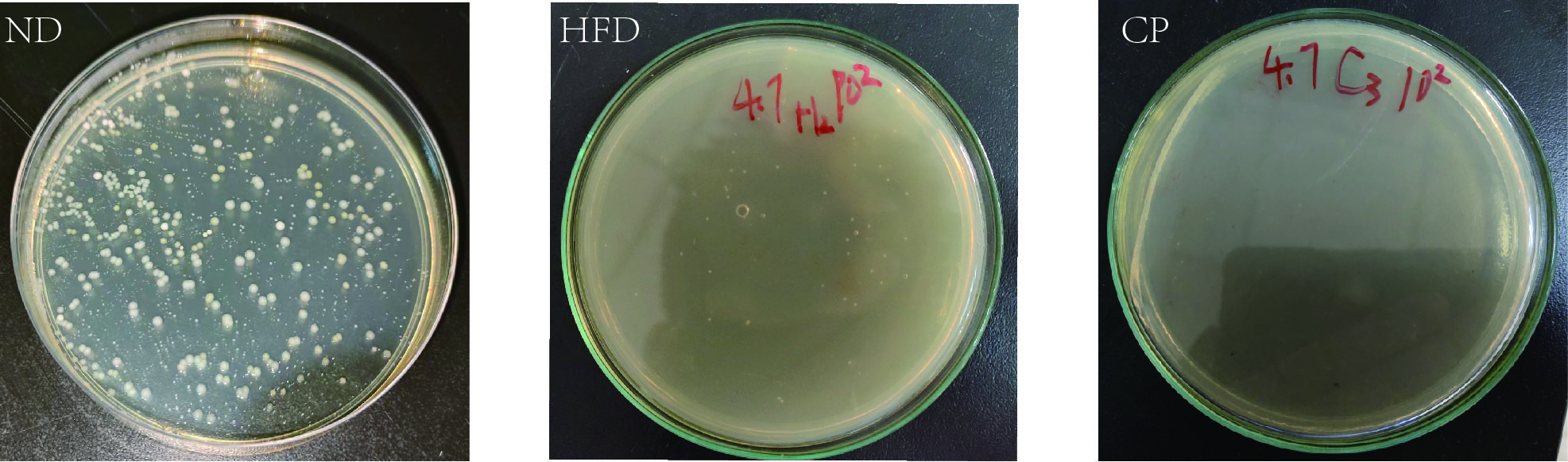
