## Supplementary material for "*Cyclocarya paliurus* polysaccharides improve high fat diet-fed mice lipid dysmetabolism through gut microbiota": Table1

| Ingredient | Normal food(g) | High fat-diet food(g) |
| --- | --- | --- |
| corn | 527 | 177 |
| casein | 200 | 250 |
| sucrose | 100 | 200 |
| bran | 80 | 80 |
| maltodextrin | 35 | 35 |
| multivitamin | 10 | 10 |
| cysteine | 3 | 3 |
| lard | 20 | 220 |
| bean oil | 25 | 25 |
| Salt | 5 | 5 |

Table1：Food ingredient list per 1 kg.
