## Supplementary material for "*Cyclocarya paliurus* polysaccharides improve high fat diet-fed mice lipid dysmetabolism through gut microbiota": Table2

|  | forward | reverse |
| --- | --- | --- |
| β-Actin | 5′-GCCCTGGACTTCGAGCAAGA-3′ | 5′-TGCCAGGGTACATGGTGGTG-3′ |
| IL-1β | 5′-TCCATGAGCTTTGTACAAGGA-3′ | 5′-AGCCCATACTTTAGGAAGACA-3′ |
| IL-6 | 5′-GTTCTCTGGGAAATCGTGGA-3′ | 5′-TGTACTCCAGGTAGCTA-3′ |
| TNF-α | 5′-AGACCCTCACACTCAGATCA-3′ | 5′-TCTTTGAGATCCATGCCGTTG-3′ |
| Resistin | 5′-GCATCCTTACCCACTGAACCAT-3′ | 5′-TCAAGCCTGTCTTTCACCTCTG-3′ |

Table2：Gene primer sequence.
